# G-quadruplexes regulate miRNA biogenesis in live zebrafish embryos

**DOI:** 10.1101/2023.01.30.526225

**Authors:** Tomás J. Steeman, Andrea M. J. Weiner, Aldana P. David, Andrés Binolfi, Nora B. Calcaterra, Pablo Armas

**Affiliations:** Instituto de Biología Molecular y Celular de Rosario (IBR), Consejo Nacional de Investigaciones Científicas y Técnicas (CONICET) - Universidad Nacional de Rosario (UNR), Ocampo y Esmeralda, (S2000EZP) Rosario, Argentina; Plataforma Argentina de Biología Estructural y Metabolómica (PLABEM), Ocampo y Esmeralda, (S2000EZP) Rosario, Argentina

**Keywords:** miR-150, G-quadruplex, microRNA, zebrafish, embryonic development

## Abstract

RNA guanine quadruplexes (G4s) regulate RNA functions, metabolism, and processing. G4s formed within precursors of microRNAs (pre-miRNAs) may impair pre-miRNAs maturation by Dicer, thus repressing mature miRNA biogenesis. As miRNAs are essential for proper embryonic development, we studied the role of G4s on miRNA biogenesis *in vivo* during zebrafish embryogenesis. We performed a computational analysis on zebrafish pre-miRNAs to find putative G4 forming sequences (PQSs). The precursor of the miRNA 150 (pre-miR-150) was found to contain an evolutionarily conserved PQS formed by three G-tetrads and able to fold *in vitro* as G4. MiR-150 controls the expression of *myb*, which shows a well-defined knock-down phenotype in zebrafish developing embryos. We microinjected zebrafish embryos with *in vitro* transcribed pre-miR-150 synthesized using either GTP (G-pre-miR-150) or 7-Deaza-GTP, a GTP analogue unable to form G4s (7DG-pre-miR-150). Compared to embryos injected with G-pre-miR-150, embryos injected with 7DG-pre-miR-150 showed higher levels of miRNA 150 (miR-150) and lower levels of *myb* mRNA and stronger phenotypes associated with *myb* knock-down. The incubation of pre-miR-150 prior to the injection with the G4 stabilizing ligand pyridostatin (PDS) reverted gene expression variations and rescued the phenotypes related to *myb* knock-down. Overall, results suggest that the G4 formed in pre-miR-150 functions *in vivo* as a conserved regulatory structure competing with the stem-loop structure necessary for miRNA biogenesis.

## 1. Introduction

Micro RNAs (miRNAs) are the most studied small noncoding RNAs (ncRNAs) due to their pivotal function in gene expression control, regulating up to 60% of the human protein-coding genes by repressing translation, promoting degradation of the target mRNA, or enhancing translation at the post-transcriptional level [1]. MiRNAs are single stranded RNAs of ≈ 23 nucleotides in length [1], synthesized by the transcription of longer primary miRNAs (pri-miRNAs) and subsequent multi-step maturation. In animals, pri-miRNAs adopt a long hairpin-like structure and are cleaved in the nuclei by the microprocessor complex composed of Drosha, DiGeorge critical region 8 (Dgcr8), and other proteins, rendering the precursor miRNAs (pre-miRNAs) structured as a stem-loop [2]. The pre-miRNAs are then exported to the cytoplasm, where the stem-loop is cleaved by the Dicer exonuclease, generating a short RNA duplex. One of the two strands of the duplex is degraded while the other one, the mature miRNA, is loaded onto the Ago2 protein within the RNA induced silencing complex (RISC), enabling the miRNA to bind and repress target mRNAs [3] through the induction of deadenylation of the poly(A) tail, mRNA destabilization and decay, and/or inhibition of translation [4].

The onset of various human diseases, including cancer [5,6], neurodegenerative diseases [7], and cardiovascular diseases [8], is due to aberrations in the regulatory processes involving miRNA expression and processing. Besides, miRNAs are essential for proper developmental embryogenesis, cell differentiation, organogenesis, growth, and programmed cell death [9,10]. During embryonic development, miRNAs are expressed in distinct spatial and temporal patterns with functions in the coordination of cell replication timing and cell fate transitions [11]. Mis-expression of miRNA leads to aberrant phenotypes [12].

MiRNAs biogenesis depends on structure-based mechanisms through the formation of complex secondary and tertiary structures, including bulges, hairpins, stem-loops, duplex-, triplex- and G-quadruplex-motifs, which allow interactions with both proteins and other nucleic acids [13]. Stem-loops in pre-miRNA are key structural players for miRNA processing [2]. Secondary structures competing with stem-loops can inhibit Dicer cleavage and, consequently, the biogenesis of miRNAs. Among others, G-quadruplexes (G4s) structures can mold the biogenesis and function of miRNAs [14–17]. G4s are thermodynamically stable four-stranded secondary structures that can be formed by the folding of single-stranded guanine (G)-rich DNA or RNA sequences [18]. Putative G-quadruplex sequences (PQSs) present at least four contiguous tracts of two or more guanine nucleotides interspersed with short nucleotide sequences forming the G4 loops. G4s structure is characterized by the stacking of two or more planar arranges of four Gs (called G-tetrads) stabilized by lateral Hoogsteen-type hydrogen bonds, π–π interactions between the Gs in stacked G-tetrads, and by the coordination of monovalent cations. K^+^ is considered the main intracellular G4-stabilizing cation, while Li^+^ is considered as a non-stabilizing or neutral cation in G4 folding and stability [19]. G4s were found in all the taxonomic phyla [20] and were reported involved in the regulation of the nucleic acids metabolism [18]. RNA G4s (rG4s) have been validated as regulators of transcription termination, pre-mRNA processing, mRNA targeting, mRNA translation and maintenance of telomere homeostasis [21,22].

Emerging reports have also informed the presence of rG4s in some types of ncRNAs, mainly in long ncRNAs (lncRNAs) and miRNAs biogenesis and functions [14–17]. Several reports show that rG4s play regulatory roles in every step of the miRNA biogenesis [14– 17]. A few reports suggest that formation of rG4s in pri-miRNAs impact on the processing of the pri-miRNAs by preventing Drosha-DGCR8 binding and processing, eventually suppressing the biogenesis of the pre-miRNAs and ultimately lading to lower levels of miRNAs [23–25]. More abundant evidence has been achieved for the role of rG4s formed in pre-miRNAs. All the studies indicate that rG4s in the pre-miRNAs compete with the stem-loop structures recognized by Dicer, thus inhibiting Dicer-mediated maturation and consequently reducing the mature miRNAs levels [26–33]. In addition, a recent report demonstrated that G4s bind to Dicer and inhibit its activity [34]. The formation and function of rG4s in mature miRNAs has been also reported in several works, suggesting a role in impairing miRNA binding to target mRNAs [35–37]. MiRNAs expression and functions are also mediated by G4s present in other molecules than the miRNAs or their precursors: e.g. DNA G4s were reported regulating the transcription of miRNA genes [38], and rG4s in mRNAs (mainly in 3’ UTRs) were described as regulators of miRNAs accessibility to their target sequences [39–42].

In recent years the knowledge about the biological role of the rG4s in miRNAs biogenesis and functions has made significant progress. However, most of the data have been achieved by bioinformatic predictions, *in vitro* structural and biochemical assays, or by performing a few assays in cultured cells, but only a couple of works have assayed the function of rG4s in the miRNA biology *in vivo* using live, complex, and multicellular organisms. One of these works reported the presence of a conserved rG4 within the pri-miRNA of miR-23b/27b/24-1 cluster, which prevents *in vivo* the processing by Drosha-Dgcr8 and thus suppresses the biogenesis of the three miRNAs involved in regulation of cardiac function in rats [25]. The other work identified an rG4 in the pre-miR-26a-1 that impairs pre-miR-26a-1 maturation, resulting in a decrease in the miR-26a levels. This leads to an increase in the miR-26a targets important for hepatic insulin sensitivity and lipid metabolism in mice [32].

We focused our study on the role of rG4s in the biogenesis of miRNAs during vertebrate embryonic development. During embryonic development, gene expression is orchestrated by specific and highly evolutionarily conserved mechanisms that take place accurately, both at spatial and temporal levels. The alteration of the fine-tuning of gene expression at any of the different developmental stages may set up specific and well defined phenotypes [43]. In this context, we wondered whether rG4s may contribute to the regulation of miRNAs functions that control genes required for the proper vertebrate embryonic development. We used zebrafish (*Danio rerio*), a vertebrate model ideal for studying embryonic development that shows high genetic similarity with humans [44], and that contains conserved miRNAs in its genome [12,45].

In this work, we present evidence gathered by using combined computational and experimental analyses showing that an evolutionarily conserved rG4 found in the precursor of miRNA 150 regulates *in vivo* the miRNA 150 biogenesis, thus modulating the expression of the specific target gene *myb* during zebrafish embryonic development. To our knowledge, this is the first work reporting the function of an rG4 as a regulatory switch to fine-tune the biogenesis of a miRNA during vertebrate embryonic development.

## 2. Results

### 2.1. Bioinformatic identification of pre-miRNAs containing evolutionarily conserved PQSs

We performed a computational analysis on zebrafish pre-miRNAs to search for PQSs within the sequences of zebrafish pre-miRNAs (Figure 1a). A total of 1114 zebrafish sequences predicted as pre-miRNAs were downloaded from miRBase [46] and Ensembl [47] databases (346 sequences, Supplementary Table S1, and 768 sequences, Supplementary Table S2, respectively). Only 350 of the downloaded sequences were annotated and associated with an identified miRNA name (the 346 from miRBase and 4 additional from Ensembl). Then, we used the downloaded sequences to search for PQSs displaying the canonical consensus G_2+_N_1-7_G_2+_N_1-7_G_2+_N_1-7_G_2+_; i. e., (G4s containing two or more G-tetrads, by using Quadparser program [22]. We found that 174 sequences (33 from miRBase, and 141 from Ensembl) contained two-G-tetrads PQSs (Supplementary Table S3). Among them, ≈ 52% of the sequences from miRBase (17/33) and ≈ 19% of the sequences from Ensembl (27/141) contained PQSs predicted with high probability to form G4s (Supplementary Table S3). Four sequences containing the canonical three G-tetrads PQSs (G_3+_N_1-7_G_3+_N_1-7_G_3+_N_1-7_G_3+_) were found, all of them displaying high probability to form G4s (Supplementary Table S4). Three of them were only retrieved from the Ensembl database and had no associated miRNA name, while the fourth was retrieved from both databases and was annotated as the precursor of miRNA 150 (pre-miR-150). We focused our study on the PQS found in pre-miR-150 sequence as a potential regulator of the miRNA 150 (miR-150) biogenesis mainly because it showed the highest scores in all the rG4 predictors (Supplementary Table S3 and S4). miR-150 is a hematopoietic cell-specific miRNA with essential regulatory roles in both normal and malignant hematopoiesis, becoming a relevant potential therapeutic target in treating various types of hematopoietic malignancies [48], and is involved in a variety of solid tumors, including breast, lung, and gastric cancer [49]. One of the best described and conserved targets of miR-150 is *myb* (also known as *c-myb*), a proto-oncogene encoding a transcription factor involved in the proliferation, differentiation, and survival of hematopoietic progenitors as well as leukemia and certain solid tumors [48,49]. Mis-regulation of *myb* causes well-defined phenotypes in developing zebrafish [50].

**Figure 1.**
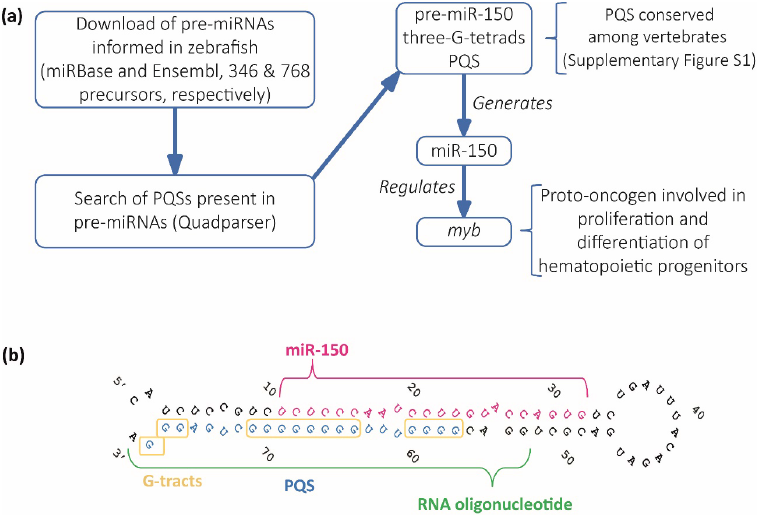
Identification and selection of pre-miRNAs containing PQSs. **(a)** Diagram of the computational search strategy of zebrafish pre-miRNAs sequences containing PQSs. **(b)** Sequence and stem-loop structure of zebrafish pre-miR-150. The mature miR-150 sequence (pink), the PQS (blue), the synthetic RNA oligonucleotide used in subsequent experiments (green), and the G-tracts (yellow boxes) are detailed.

The PQS found in zebrafish pre-miR-150 is located at the 3’ end in the stem of the predicted stem-loop structure, and is partially complementary to the mature miR-150 sequence (Figure 1b). Therefore, the formation of a G4 might interfere with the formation of the stem-loop structure, thus potentially impeding the pre-miR-150 processing by Dicer. The alignment of the sequences of pre-miR-150 orthologues showed that miR-150 sequence is highly conserved and is found only in vertebrates (Supplementary Figure S1). All the assessed pre-miR-150 sequences contain PQSs located at the 3’ ends, which do not overlap with the mature miRNA-150 sequences (Supplementary Figure S1 and Supplementary Table S5). This finding suggests a functional evolutionary conservation of the G4 forming sequences and a role in miR-150 biogenesis. In agreement, a previous computational prediction found that *Danio rerio* pre-miR-150 (dre-pre-miR-150) was the unique pre-miRNA containing the PQS identified here, responding to an extended canonical PQS with four tracts of three consecutive guanines and loops ranging from 1 to 12 nucleotides (G_3+_N_1-12_G_3+_N_1-12_G_3+_N_1-12_G_3+_) [28].

### 2.2. In vitro characterization of the G4 formed by the PQS present in zebrafish pre-miR150

The formation of G4 by the PQS of human pre-miR-150 (hsa-pre-miR-150) was studied *in vitro* as a putative probe for the detection of Nucleolin in liquid biopsies for lung cancer diagnosis, prognosis, and patient response. This work reported the formation of a stable parallel G4 in the presence of KCl or when complexed with the G4 ligand PhenDC3 [51]. However, the formation of the G4 in dre-miR-150 was not previously explored, nor was the function of the G4 in regulating the pre-miR-150 processing. Here, we used a synthetic RNA oligoribonucleotide (Figure 1b and Supplementary Table S6) to assay *in vitro* G4 formation by the PQS identified in the dre-pre-miR-150 using four different spectroscopic approaches: Circular Dichroism (CD) Spectroscopy (Figure 2a), Thermal Difference Spectroscopy (TDS) (Figure 2b), Thioflavin T (ThT) fluorescence (Figure 2c), and 1 dimension (1D) ^1^H Nuclear Magnetic Resonance (NMR) (Figure 2d). The CD spectra show the typical pattern of peaks associated with the parallel G4 structure, showing an increase of a positive peak around 264 nm and a negative peak around 240 nm in response to the presence of increasing concentrations of K^+^ (Figure 2a). The CD spectra observed in the presence of Li^+^ did not increase the characteristic G4 peaks. Thermal stability calculated by CD melting (Supplementary Figure S2) showed that the G4 is highly stable in the presence of 100 mM K^+^, since melting temperature (Tm) could not be estimated due to incomplete melting, and an estimated Tm of 59.5 °C was observed for the G4 folded in the presence of 1 mM K^+^. In addition, TDS spectra showed the typical G4 signature with two positive peaks around 243 and 273 nm and a negative peak at 295 nm (Figure 2b), and ThT fluorescence assays showed that the folded PQS markedly enhance the ThT fluorescence near 70-fold (Figure 2c). Consistent with these results, 1D ^1^H NMR showed defined signals around 11–12 ppm (Figure 2d), confirming the presence of Hoogsteen bonds and G4 structures. Moreover, the absence of signals at 13 and 15 ppm indicate that there was no significant formation of stable Watson-Crick or i-moif structures [52].

**Figure 2.**
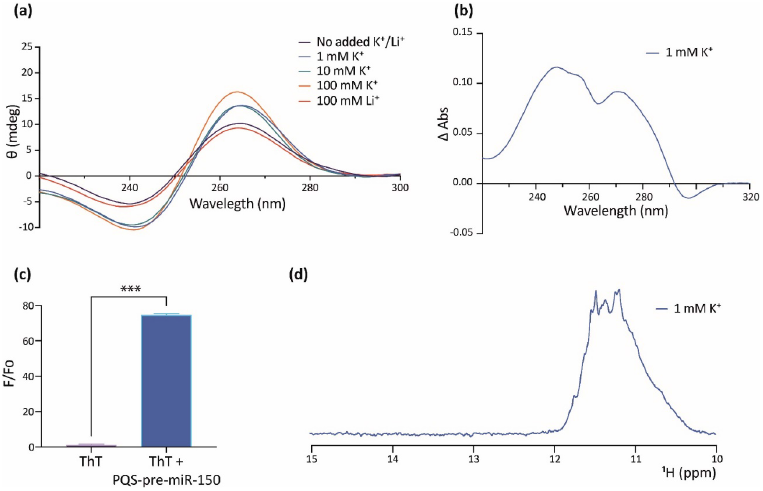
*In vitro* evidence of the formation of a G4 structure in the PQS present in pre-miR-150. **(a)** CD spectroscopy of the folded RNA oligonucleotide containing the pre-miR-150 PQS in different K^+^ and Li^+^ concentrations. **(b)** TDS spectrum of the RNA oligonucleotide containing the pre-miR-150 PQS (70°C spectrum – 20°C spectrum) folded in the presence of 1 mM K^+^. **(c)** ThT fluorescence assay with the RNA oligonucleotide containing the pre-miR-150 PQS folded in the presence of 1 mM K^+^. Two-tailed *t*-Student, p < 0.0001. **(d)** Imino proton region of the 1D ^1^H NMR spectrum of the RNA oligonucleotide containing the pre-miR-150 PQS folded in the presence of 1 mM K^+^.

These results indicate that the PQS found in dre-pre-miR-150 folds *in vitro* as a highly stable G4.

### 2.3. Analysis of pre-miR150 G4 as a regulatory element in developing zebrafish embryos

The miR-150/*myb* regulatory pathway has been described in zebrafish, for which specific phenotypes in developing embryos overexpressing miR-150 or knocked-down in *myb* expression were reported [50,53]. Overexpression of hsa-miR-150 in zebrafish embryos caused a reduction of *myb* mRNA levels in 24 hours post-fertilization (hpf) staged embryos and distinctly abnormal phenotypes in 48 hpf staged larvae, with the most obvious and common phenotypes including shortened trunk and reduced eye sizes, together with a slower heartbeat, and sluggish blood flow. Similar abnormal phenotypes were observed when *myb* expression was knocked-down by microinjecting specific morpholino. These reports make miR-150 an interesting case for studying the regulation of miRNAs biogenesis and function by G4s during embryonic development.

So, we studied the role of the G4 present in the dre-pre-miR-150 in living zebrafish embryos by microinjecting dre-pre-miR-150 into one-cell staged zygotes. Transcripts containing dre-pre-miR-150 sequence were synthesized *in vitro* using GTP in the ribonucleotide mixture (obtaining G-pre-miR-150) or alternatively the GTP analog 7-Deaza-GTP (obtaining 7DG-pre-miR-150). 7-Deaza-GTP prevents Hoogsteen base-pairing and G4 formation but does not affect the formation of Watson–Crick base-pairing [54], thus allowing only the formation of the stem-loop structure (Figure 3a). Injection of G-pre-miR-150 caused a significant increase of miR-150 levels at 24 hpf staged embryos (Figure 3b), which was dose-dependent to the amount of injected pre-miR-150 (Supplementary Figure S3a). Notably, injection of 7DG-pre-miR-150 caused a significantly higher increase of miR-150 levels (even significantly higher than those observed for G-pre-miR-150 overexpression, Figure 3b), which was also dose-dependent to the amount of injected pre-miR-150 (Supplementary Figure S3a). These results suggest that a higher proportion of the stem-loop structure is recognized by Dicer in 7DG-pre-miR-150 than in G-pre-miR-150, likely as a consequence of the absence of G4 structure in the first one. Injection of either G-pre-miR-150 or 7DG-pre-miR-150 caused a significant decrease of *myb* mRNA (Figure 3c), which was also dose-dependent to the amount of pre-miR-150 injected (Supplementary Figure S3b). To discard unspecific effects of 7-Deaza-GTP on pre-miRNA processing by Dicer, we injected 7DG- and G-pre-miR-133a, a pre-miRNA that does not contain PQSs in its sequence. No differences were observed in the miR-133a overexpression levels (Figure 3d), indicating that both transcripts were equally processed by Dicer. Moreover, injection of 7DG- and G-pre-miR-133a did not significantly alter the levels of miR-150 (Figure 3b) nor of *myb* mRNA (Figure 3c).

**Figure 3.**
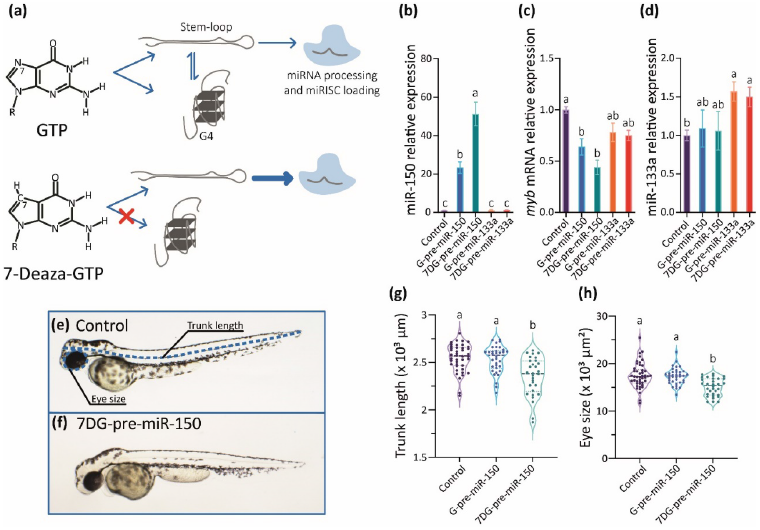
*In vivo* analysis of the role of the G4 present in pre-miR-150 during zebrafish embryo development. **(a)** Diagram of the strategy used to disrupt the formation of the G4 in the pre-miR-150 microinjected in zebrafish embryos. Reverse transcription followed by quantitative PCR (RT-qPCR) of miR-150 **(b)**, *myb* **(c)**, and miR-133a **(d)** using RNA samples from 24 hpf staged embryos microinjected with miR-150 or miR-133a precursors synthesized using either GTP (G-pre-miR-150 and G-pre-miR-133a) or 7-Deaza-GTP (7DG-pre-miR-150 and 7DG-pre-miR-133a). Ordinary 1-way ANOVA with Tukey multiple comparisons test, p < 0.005. Micrography of 48 hpf staged larvae (lateral view, head to the left) microinjected with the RNA transcribed from pSP64T+dsRED plasmid containing no cloned pre-miRNA (control, **e**) and 7DG-pre-miR-150 **(f)**, with the measured parameters indicated by blue, dashed lines in (e). Trunk length **(g)** and eye size **(h)** measurements of 48 hpf staged larvae microinjected with G-pre-miR-150 or 7DG-pre-miR-150. Ordinary 1-way ANOVA with Tukey multiple comparisons test, p < 0.001.

Then we analyzed the phenotypes of 48 hpf staged larvae injected 7DG-pre-miR-150 and G-pre-miR-150, and observed that both the trunk length and eye size were significantly reduced only in larvae injected with 7DG-pre-miR-150 (Figures 3e-h) in the highest amount injected (400 pg, Supplementary Figure S3c-d). This may indicate that the characteristic phenotypes associated with miR-150 overexpression and *myb* mRNA knock-down need threshold levels of miR-150 overexpression. This fact reinforces the notion that 7DG-pre-miR-150 produces a higher miR-150 overexpression due to a higher proportion of stem-loop and no competing G4 structure.

Data gathered so far show that the impairment of the formation of the G4 in the pre-miR-150 favors miR-150 biogenesis. Therefore, we wondered if the stabilization of the G4 in the pre-miR-150 has an opposite effect. Therefore, we tested the effect of pyridostatin (PDS) on the pre-miR-150 processing. PDS is a widely used G4 stabilizer molecule with high binding affinity to nuclear DNA G4s and cytoplasmic RNA G4s [55,56]. First, we tested *in vitro* the capability of PDS to stabilize the pre-miR-150 G4. Increasing PDS concentrations caused an increase in thermal stability of the G4 formed by the PQS present in pre-miR-150 (Figure 4). A strong stabilization was observed above 5 μM concentrations of PDS, evidenced by flattened melting curves (Figure 4a), as well as high values of CD peaks at 95°C (Figure 4b-c). So, we used 5 μM PDS for incubation of G- and 7DG-pre-miR-150 previous to their injection in zebrafish embryos. Pre-incubation of G-pre-miR-150 with PDS caused a significant reduction in miR-150 overexpression levels (Figure 5a), as well as a significant reduction in *myb* mRNA levels, although it was significantly smaller than in the absence of PDS (Figure 5b). As expected, pre-incubation of 7DG-pre-miR-150 with PDS had no effect in miR-150 overexpression levels (Figure 5a), nor in the depletion of *myb* mRNA levels (Figure 5b). These results are consistent with the stabilizing effect of PDS on the G4 formed in G-pre-miR-150, which reduces both the stem-loop structure folding and the efficiency of Dicer processing, leading to lower mature miR-150 levels and eventually causing lower repression of *myb* expression. On the contrary, no effect of PDS was observed on 7DG-pre-miR-150, probably due to the absence of a G4 structure. In agreement, the developmental phenotypes observed after G-pre-miR-150 overexpression were reverted in the case of trunk length (Figure 5c) and partially reverted in the case of eye size (Figure 5d) by the pre-incubation of G-pre-miR-150 with PDS. Instead, no phenotypic reversion was observed in embryos injected with 7DG-pre-miR-150 pre-incubated with PDS.

**Figure 4.**
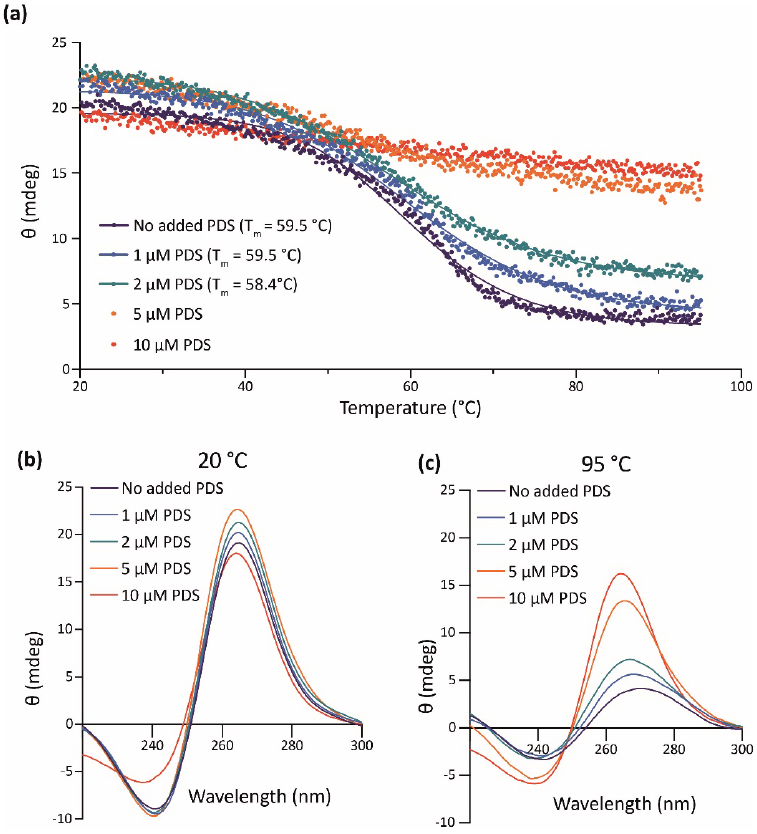
*In vitro* effect of PDS on pre-miR-150 G4 stability. **(a)** CD melting curves of the RNA oligonucleotide containing the pre-miR-150 PQS in presence of 1 mM K^+^ and different PDS concentrations. The Tm for 5 and 10 μM PDS could not be calculated due to the absence of a clear melting transition. CD spectra of the RNA oligonucleotide containing the pre-miR-150 PQS folded in presence of 1 mM K^+^ and incubated with different PDS concentrations at the start (**b**, 20°C) and at the end (**c**, 95°C) of the melting experiments.

**Figure 5.**
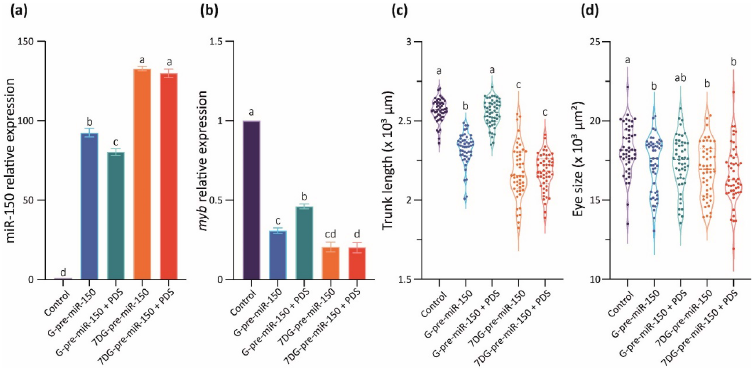
*In vivo* effect of PDS on pre-miR-150 G4 during zebrafish embryo development. RT-qPCR quantification of miR-150 **(a)** and *myb* **(b)** of 24 hpf staged embryos microinjected with G- or 7DG-miR-150 in the absence or in the presence of 5 μM PDS. Ordinary 1-way ANOVA with Tukey multiple comparisons test, p < 0.001. Trunk length **(d)** and eye size **(d)** measurements of microinjected 48 hpf staged larvae microinjected with G- or 7DG-miR-150 in the absence or in the presence of 5 μM PDS. Ordinary 1-way ANOVA with Tukey multiple comparisons test, p < 0.001.

Here we used two different strategies, one inhibiting G4 formation by using 7-Deaza-GTP for pre-miRNA synthesis and the other promoting G4 formation by the incubation of miRNA with PDS prior to embryos injection, to show that the G4 formed by pre-miR-150 functions *in vivo* as a structural switch preventing the folding of the stem-loop structure and the miR-150 biogenesis.

## 3. Discussion

Here we focused our interest in pre-miR-150 that was the only zebrafish pre-miRNA annotated in miRBase containing a three-G-tetrads PQS and displaying the highest scores in G4RNAscreener, a robust bioinformatic tool developed for identification of RNA PQSs that combines three different non-motif-based G4 predictors [57]. However, in the search for PQS capable of forming rG4 with possible regulatory roles in zebrafish miRNAs, we observed that several pre-miRNAs (33/346, ≈ 10%) annotated in miRBase contain two-G-tetrads PQS, being ≈ 52% of them (17/33, ≈ 5% of the total) predicted with a high probability of forming G4 by G4RNAscreener (Supplementary Table S3). In agreement, G4RNAscreener predicted that 2% of human pre-miRNA contains PQSs, all of them overlapping with a processing site [58]. The G4s folded in RNA are intrinsically more stable than those formed in DNA [59] and can be stable enough to be considered as structures formed transiently in the cellular context. rG4s fit perfectly with the intrinsic nature of RNA molecules as a flexible molecule that undergoes dynamic changes between conformational states in response to environmental and physical factors (interaction with proteins, ions, temperature changes, ligands and/or other nucleic acids), thus exerting their function through a structure-based mechanisms [14,60]. Therefore, although the two-G-tetrads G4s have been reported as less stable than G4s containing three or more G-tetrads [61], pre-miRNAs containing two-G-tetrads PQSs with high G4RNAscreener scores are interesting for further studies.

The inspection of pre-miR-150 orthologues sequences showed that miR-150 sequence is highly conserved and suggests that it is exclusive of vertebrate species (38 available sequences in miRBase). The analysis of the presence of PQSs within these pre-miR-150 orthologues revealed that, although the PQS is less conserved in sequence identity than the miRNA sequence, all the pre-miR-150 orthologues contain PQSs within the 3’ ends and in regions not overlapping with the mature miR-150. This suggest a functional conservation of the G4 as a structure able to compete with the formation of the stem-loop and with the processing by Dicer. Moreover, the *myb* oncogene is a conserved target of miR-150 [50], thus making the regulatory mechanism of miR-150 biogenesis described here also feasible in other organisms and/or other processes involving *myb* oncogene.

Here, the repressor function of the rG4 found in pre-miR-150 on the biogenesis of miR-150 was demonstrated by the overexpression the pre-miR-150 *in vivo* in developing zebrafish embryos. Two different approaches were used, one inhibiting and another promoting the formation of the G4 in the pre-miR-150. To inhibit the G4 formation, we used the GTP analog 7-Deaza-GTP for preventing Hoogsteen base-pairing and G4 formation, while allowing the formation of Watson–Crick base-pairing. A similar strategy was previously used for the study of the role of rG4s in the processing of pre-mRNAs during splicing, evidencing the structural function of rG4s co-existing with other competing secondary structures [54,62]. An alternative strategy for preventing G4 formation is to mutate specific Gs in the G-tracts of the PQS. However, the alteration of the PQS may in turn alter the interaction of the pre-miRNA with other biomolecules, such as proteins or other nucleic acids. In the particular case of pre-miR150, since the PQS is located in the stem of the stem-loop structure (Figure 1b), mutations of the PQS may also affect the stem-loop folding, which may require compensatory mutations in the complementary strand to maintain the structural moiety. Moreover, as the PQS is complementary with the mature miRNA sequence (Figure 1b), compensatory mutations may change the mature miRNA sequence, with consequences on the recognition of targets. Therefore, the use of 7DG-pre-miR-150 for overexpression and the comparison with results using G-pre-miR-150, turn into a robust and elegant strategy for evidencing the effect of the rG4 in the pre-miRNA processing, avoiding mutagenesis strategies. On the other hand, to promote the G4 formation, we incubated the pre-miRNA with PDS, a widely used G4 stabilizer molecule with high binding affinity to most DNA and RNA G4s [55,56]. So, the strategy of the incubation of the *in vitro* synthesized pre-miRNAs with PDS previous to the microinjection in zebrafish embryos reduced the chances of PDS to interact with other DNA and RNA G4s likely present in embryonic cells, allowing to target the effect of the ligand mainly to the injected pre-miRNA molecule. The two strategies used here represent original approaches, since most of the assays formerly used to assess the function of rG4s in pre-miRNAs processing were performed in cultured cells or *in vivo* by using classical approaches of mutagenesis and incubation of cells with G4-stabilising or G4-destabilising ligands.

Our results indicate that the evolutionarily conserved rG4 formed in the pre-miR-150 may function *in vivo* as a conserved regulatory structure competing with the stem-loop structure necessary for miR-150 biogenesis, thus regulating its levels and the expression of its target genes These evidences support the idea that rG4s in pre-miRNAs exist in a dynamic equilibrium with the stem-loop structure, which may be sensitive to subtle changes in the intracellular conditions and, ultimately allowing proper regulation of gene expression. Moreover, our results contribute to the knowledge about miRNA expression control and allow to delineate novel strategies to regulate miRNA function by targeting rG4s with specific ligands, antisense strategies or intervening the action of proteins that bind and stabilize or unfold rG4s.

## 4. Materials and Methods

### 4.1. Bioinformatics

The sequences of zebrafish (*Danio rerio*) pre-miRNAs were downloaded from the Ensemble (www.ensembl.org, Release: 89, Assembly: GRCz10) [47] and miRBase (www.mirbase.org) [46] databases. Full names, ID numbers and sequences are shown in Supplementary Tables S1 and S2. Putative G4 sequences (PQSs) were identified with Quadparser [22], searching for the consensus sequence G_2+_N_1-7_G_2+_N_1-7_G_2+_N_1-7_G_2+_.

### 4.2. Oligonucleotides

For the spectroscopic studies, a synthetic single-stranded desalted oligoribonucleotide (pre-miR-150-PQS, ST1) was purchased from Invitrogen (Carlsbad, CA, USA) resuspended in nuclease-free water and stored at -20°C until use. Concentration was determined by absorbance at 260 nm (NanoVue Plus, Biochrom, Holliston, MA, USA) using the molar extinction coefficient provided by the manufacturer. For all experiments, the oligoribonucleotide was diluted to the final experimental concentration in 10 mM Tris-HCl pH 7.5 containing varying KCl or LiCl concentrations, as indicated in each figure, and folded by heating for 5 min at 95°C and slowly cooling to 20°C. For cloning and RT-qPCR experiments, DNA oligonucleotides were purchased from Macrogen (Seoul, South Korea), resuspended in nuclease-free water, and stored at -20°C until use. All sequences are shown in Supplementary Table S6.

### 4.3. Circular dichroism (CD) spectroscopy

Circular dichroism spectra (ellipticity, Ө) were acquired at 20°C between 220-300 nm using a Jasco-1500 spectropolarimeter (10 mm quartz cell, 100 nm/min scanning speed, 1 s response time, 1 nm data pitch, 2 nm bandwidth, average of four scans), as described elsewhere [63]. An oligoribonucleotide concentration of 2 μM was used, in the absence or in the presence of varying KCl (1, 10, or 100 mM), or 100 mM LiCl. The spectral contribution of buffers, salts, and ligands was appropriately subtracted. The CD melting curves were recorded by ellipticity measurements between 20 and 95°C, at the wavelength corresponding to the maximum observed at 20°C for the positive peak around 264 nm, as described elsewhere [63]. For melting temperature (Tm) calculation, data was analyzed with Prism 9 (GraphPad) with a non-linear least square fitting procedure assuming a two-state transition of a monomer from a folded to an unfolded state with no change in heat capacity upon unfolding. For ligand experiments, PDS (pyridostatin trifluoroacetate salt, Sigma-Aldrich SML0678) was added at different concentrations (0, 1, 2, 5, and 10 μM) after oligoribonucleotide folding and incubated for 30 min at 37°C before CD spectra and melting recording.

### 4.4. 1D ^1^H Nuclear magnetic resonance (NMR)

NMR spectra were acquired at 20°C on a 700MHz Bruker Avance III spectrometer (Bruker Biospin, MA, USA) equipped with a triple resonance inverse NMR probe (5 mm1H/D-13C/15N TXI). Experiments were performed on 50 μM RNA oligonucleotide samples folded in the presence of 1 mM KCl. 1D ^1^H NMR spectra were acquired using the zgesgp pulse sequence for efficient water suppression [64]. We used 8K points, 1024 scans, a recycling delay of 1.4 s and a sweep width of 22 ppm. Experimental time for each NMR spectrum was 29 min. Spectra were processed by exponential multiplication (line broadening of 10 Hz) and baseline correction. NMR acquisition, processing, and analysis was done using Topspin 3.5 (Bruker, Biospin, MA, USA).

### 4.5. Thermal Difference Spectroscopy (TDS)

The oligoribonucleotide folded at 2 μM in the absence or in the presence of 1 mM KCl or 1 mM LiCl was scanned to measure absorbance from 220 to 320 nm using a scan speed of 100 nm/min and a data interval of 1 nm in a 10 mm quartz cell. Spectra were measured at 20 and 70°C using a Jasco V-630BIO spectrophotometer with Peltier temperature control. The difference between the spectra at these temperatures (Abs 70°C – Abs 20°C) was calculated and plotted to obtain the TDS as previously described [63].

### 4.6. Thioflavin T (ThT) fluorescence assays

ThT (Sigma-Aldrich T3516) fluorescence assays were performed as previously described [65]. Fluorescence emission measurements were performed using a microplate reader (Synergy 2 MultiMode Microplate Reader, BioTek, VT, USA) with an excitation filter of 485 ± 20 nm and an emission filter of 528 ± 20 nm. Each sample was tested in triplicate and fluorescence values were relativized to ThT fluorescence in absence of oligonucleotides (F0). A threshold of a 10-fold increase was used for considering G4 formation.

### 4.7. Animal care

Zebrafish specimens were handled according to relevant national and international guidelines and ethically authorized by the Internal Committee for the Care and Use of Laboratory Animals of the Facultad de Ciencias Bioquímicas y Farmacéuticas - Universidad Nacional de Rosario, which has been accepted by the Ministerio de Salud de la Nación Argentina (expedient 6060/374, resolution 207/2018). Adult zebrafish were maintained at 28°C on a 14-10 h light-dark cycle as previously described [66]. Mating was carried out by crossing three males with four females in the same spawning tank. Embryos were staged according to morphological development in hours or days post-fertilization at 28°C as described elsewhere [67].

### 4.8. Pre-miR overexpression by microinjection

For the overexpression experiments, the genomic regions of zebrafish pre-miR-150 (miRBase accession code MI0002016; chr3: 32571216-32571517) and pre-miR-133a (miRBase accession code MI0001993; chr2: 4113916-4114439) were amplified by PCR and cloned into a pSP64T+dsRED vector [68], using EcoRI and XhoI restriction sites included in the primers (Supplementary Table S6). The plasmid containing no cloned pre-miRNA was used as control. RNA for microinjection was *in vitro* synthesized using as a template the plasmids previously linearized with BamHI (Promega) and transcribed using mMESSAGE mMACHINE SP6 kit (Invitrogen). The NTP-CAP mix included with the transcription kit was replaced with a custom mix containing CAP 4 mM, ATP, CTP, UTP, and either GTP or 7-deaza-GTP 10mM each (7-Deazaguanosine-5’-Triphosphate, TriLink, San Diego, CA, USA). One-cell staged embryos were microinjected with 200 and 400 pg of the transcripts and incubated at 28°C until they reached the stage for collection indicated for each experiment. PDS was added at a final concentration of 5 μM to the transcripts and incubated for 30 min at 25°C before microinjection.

### 4.9. Reverse transcription followed by quantitative PCR (RT-qPCR)

For the RT-qPCR experiments, 24 hours post-fertilization (hpf) staged embryos were collected for each treatment and frozen in liquid nitrogen, and used immediately or stored at -80°C. RNA was extracted by TRIzol-Chloroform extraction (Invitrogen) according to the manufacturer’s instructions, followed by isopropyl-alcohol precipitation and pellet resuspension in nuclease-free water. RNA concentration was assessed by absorbance at 260 nm (Nanodrop). Reverse transcription was performed with M-MLV RT Enzyme (Promega, Madison, WI, USA) and oligo-dT (T12 and T16 mix) primers (Supplementary Table S6). Quantitative PCR was done with HOT FIREPol EvaGreen qPCR Mix Plus (Solis Biodyne, Tartu, Estonia) on RealPlex4 thermocycler (Eppendorf, Hamburg, Germany). For miRNA detection, specific stem-loop primers were added to the retrotranscription reaction, with a generic reverse and specific forward primers used in qPCR [69] (Supplementary Table S6). The amplification of zebrafish *rpl13* and *eef1a1l1* genes cDNAs was used as reference [70] and relative expressions were normalized to controls. All primer sequences are detailed in Supplementary Table S6. Data analysis was performed with REST2009 (Qiagen, Hilden, Germany) [71] and statistical analysis with GraphPad Prism 9 (see Supplementary Table S7 for p values of pairwise comparisons), following MIQE guidelines [72].

### 4.10. Phenotype observation

Zebrafish larvae were collected at 48 hpf and fixed with 4% paraformaldehyde in Phosphate Buffered Saline (PBS) pH 7.4 at 4°C overnight, washed 3 times with PBS and placed in glycerol 100%. The embryos were orientated laterally and photographed with an Olympus MVX10 stereoscopic microscope and Olympus C-60 ZOOM digital camera. Embryo length was measured by drawing a spline from the head to the end of the tail along the notochord, and eye size was by drawing a freehand selection around the eye, using FIJI software (National Institute of Health, Bethesda, MD, USA). Statistical analysis was performed with GraphPad Prism 9 (see Supplementary Table S7 for p values of pairwise comparisons).

## Supplementary Materials

Figure S1: Conservation of the PQSs in pre-miR-150 among vertebrate species; Figure S2: Thermal stability of the G4 formed by the PQSs in pre-miR-150; Figure S3: *In vivo* analysis of the role of the G4 present in pre-miR-150 during zebrafish embryo development; Table S1: Zebrafish pre-miRNA sequences downloaded from miRBase; Table S2: Zebrafish pre-miRNA sequences downloaded from Ensembl; Table S3: Zebrafish pre-miRNA PQSs predicted with QGRSmapper, containing two G-tetrads and scores predicted by G4RNAScreener; Table S4: Zebrafish pre-miRNA PQSs predicted with QGRSmapper, containing three G-tetrads and scores predicted by G4RNAScreener; Table S5: PQSs predicted with QGRSmapper in pre-miR-150 orthologs from 38 vertebrate species downloaded from miRBase and scores predicted by G4RNAScreener; Table S6: RNA and DNA oligonucleotide sequences; Table S7: P values of pairwise comparisons from Figures 3, 5 and S3.

## Author Contributions

P.A. and N.B.C. performed the conceptualization and design of the work. T.J.S. and A.D. carried on most bioinformatic analyses and in vitro experiments. A.M.J.W. and A.D. participated in cloning and initial set-up of *in vivo* experiments, while T.J.S. performed all the *in vivo* experiments. A.B. performed and analyzed NMR spectroscopies. P.A. was responsible for supervision, funding, acquisition, project administration, and obtaining of resources. A.M.J.W and N.B.C. collaborated with funding. P.A. and T.J.S. conducted the original draft writing and visualization, while P.A., T.J.S. and N.B.C. performed the writing review and editing. All authors have read and agreed to the published version of the manuscript.

## Institutional Review Board Statement

Zebrafish specimens were handled according to relevant national and international guidelines and ethically authorized by the Internal Committee for the Care and Use of Laboratory Animals of the Facultad de Ciencias Bioquímicas y Farmacéuticas - Universidad Nacional de Rosario, which has been accepted by the Ministerio de Salud de la Nación Argentina (expedient 6060/374, resolution 207/2018).

## Data Availability Statement

All the data presented in this study are available in the article and supplementary material.

### Acknowledgments

T.J.S. is a Fellow of CONICET and A.D. was a Fellow of CONICET. P.A., A.W., A.B. and N.B.C. are Career Researchers of CONICET, and P.A. and N.B.C. are Professors of UNR. We are thankful to Sebastian Graziati for excellent zebrafish husbandry. We thank Andrea Coscia and Alejandro Gago for maintenance of the NMR facility.

---

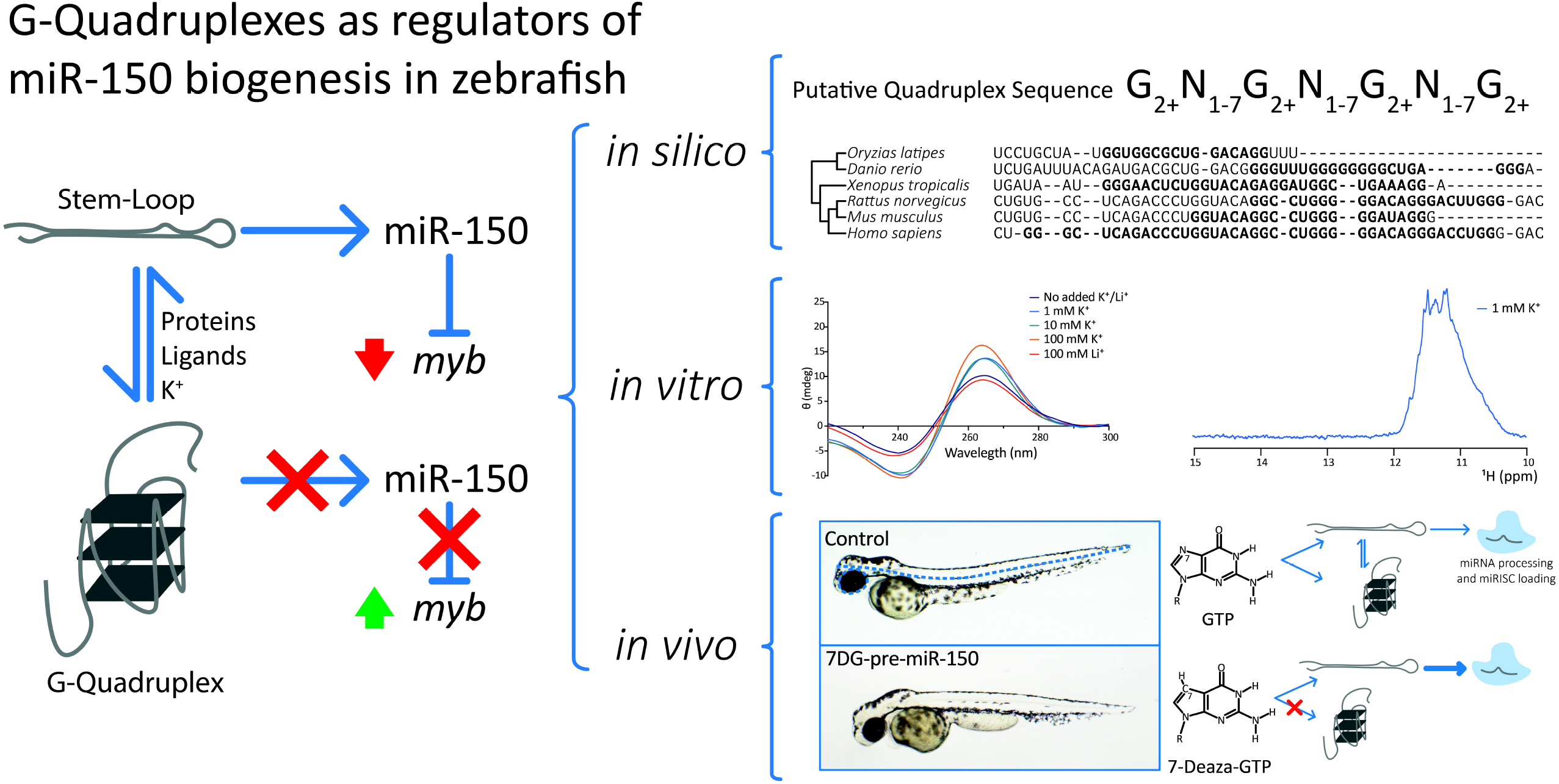

