## Supplementary Material for "G-quadruplexes regulate miRNA biogenesis in live zebrafish embryos"

**Content:**


| | Page |
|---|---|
| Supplementary Tables Description | 1 |
| Supplementary Figures | 2 |
| Supplementary Figure 1 | 2 |
| Supplementary Figure 2 | 4 |
| Supplementary Figure 3 | 4 |
| References for Supplementary Material | 5 |

#### Supplementary Tables Description

**Supplementary Table S1.** Zebrafish pre-miRNA sequences downloaded from miRBase.

**Supplementary Table S2.** Zebrafish pre-miRNA sequences downloaded from Ensembl.

**Supplementary Table S3.** Zebrafish pre-miRNA PQSs predicted with QGRSmapper, containing two G-tetrads and scores predicted by G4RNAScreener.

**Supplementary Table S4.** Zebrafish pre-miRNA PQSs predicted with QGRSmapper, containing three G-tetrads and scores predicted by G4RNAScreener.

**Supplementary Table S5.** PQSs predicted with QGRSmapper in pre-miR-150 orthologs from 38 vertebrate species downloaded from miRBase and scores predicted by G4RNAScreener.

**Supplementary Table S6.** RNA and DNA oligonucleotide sequences.

**Supplementary Table S7.** P values of pairwise comparisons from Figures 3, 5 and S3.

### Supplementary Figures

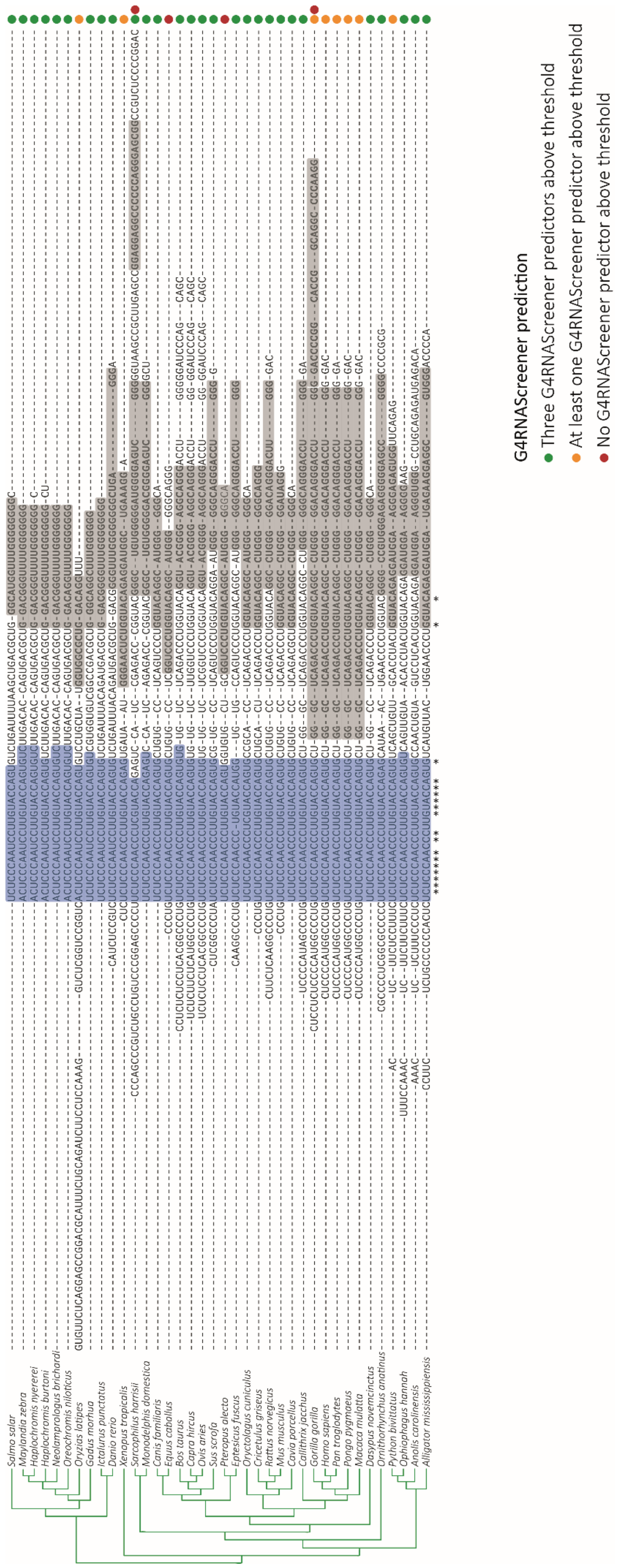


**Supplementary Figure S1**

**Supplementary Figure S1.** Conservation of the PQSs in pre-miR-150 among vertebrate species. Multiple sequence alignments were performed using Clustal Omega [1] of annotated pre-miR-150 sequences from 38 vertebrate species downloaded from miRBase [2] database. The phylogenetic tree is represented on the left. The mature miR-150 sequences are shaded in blue. PQSs found by QGRS Mapper (max length: 45, Min G-Group Size: 2, Loop size: from 1 to 15) [3] are shaded in grey. Dots on the right indicate the predicted G4 folding probability according to G4RNAScreener predictors [4] (see Supplementary Table S5 for score values). PQSs that were found with QGRS Mapper and whose scores for the three G4RNAScreener predictors were over the defined thresholds were classified with a high probability to form G4 (green dots). Those PQSs that were found with QGRS Mapper and whose score for at least one G4RNAScreener predictor was over the defined thresholds were classified with medium probability to form G4 (yellow dots). Those PQSs that were found with QGRS Mapper and whose scores for the three G4RNAScreener predictors were below the defined thresholds were classified with low probability to form G4 (red dots). *S. harrissi* and *G. gorilla* pre-miR-150 contain two predicted PQS, where the first and second probability dots correspond to the first and second PQS, respectively. The asterisks at the bottom represent 100% nucleotide conservation among all species.

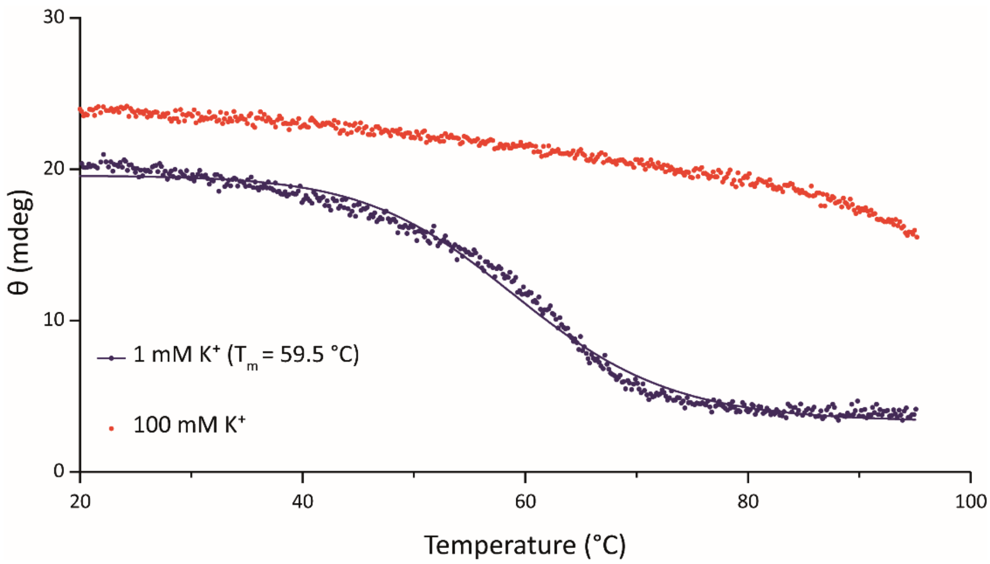


**Supplementary Figure S2.** Thermal stability of the G4 formed by the PQSs in pre-miR-150. CD melting curves of the RNA oligonucleotide containing the pre-miR-150 PQS in presence of 1 or 100 mM K+. The Tm for the G4 folded in the presence of 100 mM K+ could not be calculated due to the absence of a clear melting transition.

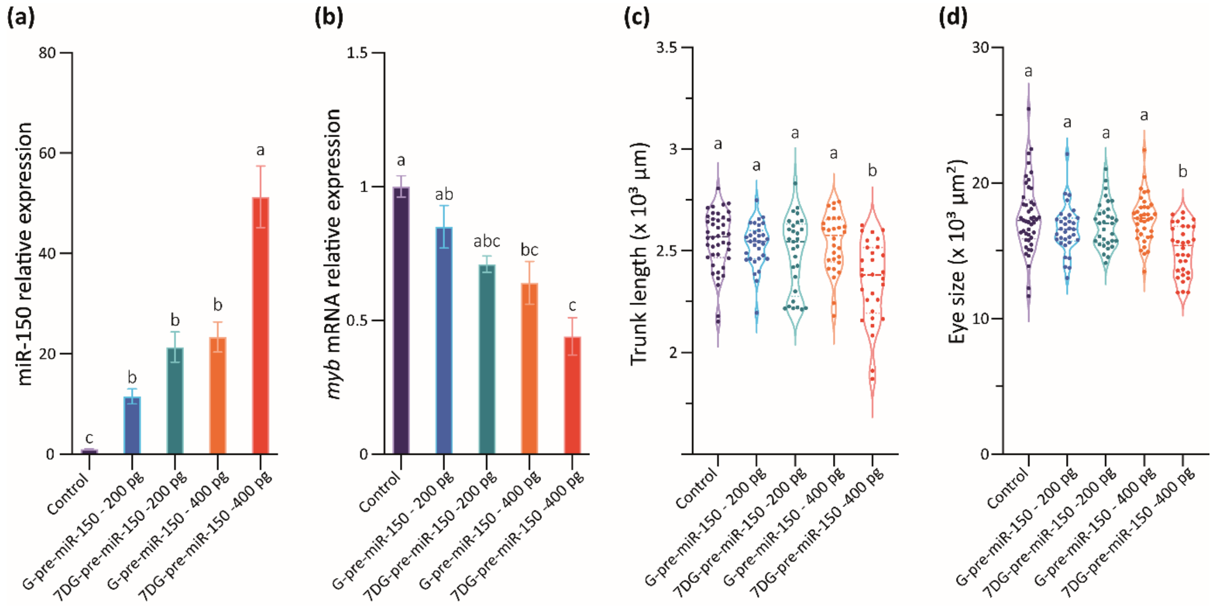


**Supplementary Figure S3.** *In vivo* analysis of the role of the G4 present in pre-miR-150 during zebrafish embryo development. RT-qPCR quantification of miR-150 **(a)** and *myb* **(b)** using RNA samples from 24 hpf staged embryos microinjected with G-pre-miR-150 or 7DG-pre-miR-150 in two different amounts (200 or 400 pg). Ordinary 1-way ANOVA with Tukey multiple comparisons test, $p < 0.005$. Trunk length **(c)** and eye size **(d)** measurements of 48 hpf staged larvae microinjected with G-pre-miR-150 or 7DG-pre-miR-150 in two different amounts (200 or 400 pg). Ordinary 1-way ANOVA with Tukey multiple comparisons test, $p < 0.001$.
